# Nutritional insensitivity to mating in male fruit flies

**DOI:** 10.1101/2023.08.11.552938

**Authors:** Mabel C Sydney, Tracey Chapman, Jennifer C Perry

## Abstract

Animals can adjust their consumption of different nutrients to adaptively match their current or expected physiological state. Changes in diet preference can arise from social and sexual experience. For example, in female *Drosophila melanogaster* fruit flies, a single mating triggers a behavioural switch in diet choice towards increased protein intake and total food consumption, which supports offspring production. In contrast, male diet choice appears to be unaffected by a single mating. However, one mating may not fully capture the impact of mating on male feeding behaviour. Males can often mate multiply in natural settings, and the costs of ejaculate production and energetic courtship may be cumulative, such that males might experience increased nutritional demands only after multiple matings. In this study we tested this idea by measuring the effect of multiple matings on the diet choice of male *D. melanogaster* fruit flies. Males were assigned to one of three mating treatments – unmated, mated once or mated five times consecutively – and then allowed to feed freely on chemically-defined diets of protein and carbohydrate. In contrast to the prediction, we found that males that mated five times did not alter the amount of food, nor the proportion of protein and carbohydrate consumed, when compared with unmated or once-mated males. This absence of a feeding response occurred despite substantial ejaculate depletion from multiple matings: males sired fewer offspring in each consecutive mating. These results reveal a lack of plasticity in male feeding behaviour according to mating status, despite substantial potential physiological costs, and highlight the remarkably distinct nutritional ecologies of males versus females.

---

Animals have complex nutritional needs, with optimal diets varying with age, sex, metabolic rate and environment (Simpson and Raubenheimer, 2012). Previous studies have demonstrated that, in many species, individuals are able to sense their own physiological state and adjust feeding to match nutritional demand, by fine-tuning consumption of micronutrients, such as vitamins and minerals, and macronutrients, including carbohydrate, protein (amino acids) and lipids (Ribeiro and Dickson, 2010; Simpson, Le Couteur and Raubenheimer, 2015). The balance of macronutrients eaten strongly affects an individual’s fitness, and the balance of protein and carbohydrate (P:C ratio) appears particularly key. For example, experiments in *Drosophila melanogaster* fruit flies show that low P:C diets can maximise the longevity of both sexes, while optimal P:C diets for reproductive success differ between males and females (Lee *et al*., 2008; Jensen *et al*., 2015; Camus *et al*., 2017; Carey *et al*., 2022; but see Reddiex *et al*., 2013). When males and females are able to choose, they favour diets that maximise reproductive success; for males this is generally a lower P:C diet than for females (Lee *et al*., 2008; Maklakov *et al*., 2008; Fanson *et al*., 2009; but see Jensen *et al*., 2015).

Mating, and the initiation of reproductive processes that stem from it, has important implications for nutrient demands in females. Females increase food intake after mating in *D. melanogaster* (Carvalho *et al*., 2006; Lee, Kim and Min, 2013; Camus *et al*., 2018) and in the two-spot ladybird *Adalia bipunctata* (Perry, 2011). Upon mating, female *D. melanogaster* markedly increase the proportion of protein and yeast in their diet, in comparison with unmated females (Barnes *et al*., 2008; Ribeiro and Dickson, 2010; Lee, Kim and Min, 2013; Jensen *et al*., 2015; Corrales-Carvajal, Faisal and Ribeiro, 2016; Camus *et al*., 2018; Newell *et al*., 2020). Macronutrient preferences are also altered during pregnancy in *Rattus norvegicus domestica* rats (Richter and Barelare, 1938; Leshner, Siegel and Collier, 1972; Simpson and Raubenheimer, 1997). In *D. melanogaster*, the increase in females’ preference for protein after mating is thought to occur to meet the demands of elevated egg production (Bownes and Blair, 1986; Drummond-Barbosa and Spradling, 2001; Lee, Kim and Min, 2013; but see Ribeiro and Dickson, 2010).

In contrast, data on the effect of mating on male dietary preference are scant. The few studies to date, conducted in *D. melanogaster* and *A. bipunctata*, report little evidence for a shift in male dietary preference after a single mating (Perry and Tse, 2013; Camus *et al*., 2018). For example, in *D. melanogaster*, a single mating had no significant effect on male diet preference (P:C ratio) or the overall quantity of food consumed (Camus *et al*., 2018). This result is in accord with the view that male ejaculates may be relatively ‘cheap’ to produce (Bateman, 1948; Trivers, 1972) and hence that males might not need to increase protein intake to replenish reserves depleted by mating. However, in species with nuptial gifts, such as the German cockroach *Blattella germanica,* there is evidence for dietary compensation (Jensen and Silverman, 2018).

It is possible that males incur cumulative costs of ejaculate production, such that costs that are insignificant after a single mating increase significantly with multiple matings. In fact, males of many species are able to mate multiply in quick succession. For example, an average *D. melanogaster* male can mate five times in four hours, with some individuals able to mate 11 times (De Crespigny, Pitt and Wedell, 2006; Douglas, Anderson and Saltz, 2020). Moreover, contrary to the ‘cheap’ male ejaculate idea, it is now understood that the production of sperm and other ejaculate components can be costly and limiting for males (Dewsbury, 1982; Olsson, Madsen and Shine, 1997; Reinhardt, Naylor and Siva-Jothy, 2011; Perry and Tse, 2013; Perry, Sirot and Wigby, 2013). For example, ejaculate components start to become depleted in male *D. melanogaster* after three sequential matings (Lefevre and Jonsson, 1962) and continue to decrease with additional matings (Hihara, 1981; Linklater *et al*., 2007; Loyau *et al*., 2012; Macartney *et al*., 2021). Male *D. melanogaster* mating five times in succession can be rendered infertile due to depletion of non-sperm ejaculate components (Hihara, 1981). Therefore, we predict that an impact of mating on male diet preference will arise after multiple matings.

To test this prediction and improve understanding of how mating impacts male diet choice, we manipulated the mating rate of male *D. melanogaster* fruit flies and then measured male diet preferences and intake. We used *D. melanogaster* because protocols for measuring its dietary preferences are well-established, as it is an important model for nutritional preference studies, and males can mate multiply. Male flies were assigned to mate once or five times consecutively, or to remain unmated. Five sequential matings approaches the average maximal daily mating rate for this species (Lefevre and Jonsson, 1962; Douglas, Anderson and Saltz, 2020). We tracked the number of viable offspring produced from each mating to confirm that the highest mating rate caused ejaculate depletion. Following the mating treatments, male diet preference was measured by offering male flies carbohydrate and protein solutions simultaneously using the Capillary Feeder (CAFE) assay (Ja *et al*., 2007). The CAFE method allows the quantification of food intake and macronutrient preference using synthetic diets with known nutritional content. We predicted that males mated at the highest rate would (1) suffer energy and ejaculate depletion, evident as cumulative reductions in their siring success with each mating bout; (2) increase total food consumption to recoup energy expenditure during courtship and mating; and (3) increase the proportion of protein eaten after mating to restore proteinaceous sperm and non-sperm components of the ejaculate.

## METHODS

### Fly stocks

Fly rearing and experiments were conducted in a 25°C humidified room under a 12-hour light-dark cycle. Experimental flies were collected from a large stock population of outbred wildtype Dahomey flies (Chapman, Trevitt and Partridge, 1994) maintained on a standard sugar-yeast-agar (SYA) diet (50g sucrose, 100g brewer’s yeast, 15g agar, 30ml Nipagin (10% solution), 3ml propionic acid, 970ml water). To generate experimental flies, eggs from the stock population were collected on grape juice-agar plates with live yeast paste. First instar larvae were transferred into glass vials containing SYA medium at a controlled density of fifty larvae per vial. Experimental adults were collected as virgins using ice anaesthesia within 4-6 hours of eclosion. Flies were housed in single sex groups of 15 males or 10 females in glass vials containing 7ml SYA medium supplemented with live yeast granules for three days before mating assays.

Experiments were carried out in four experimental blocks. In blocks one and two an additional mating treatment of three successive matings was included. Offspring data were collected from blocks one and two, dietary data were collected from blocks three and four, and mating data were collected from all four blocks.

### Mating treatment

Experimental males were randomly assigned to one of three mating treatments: zero matings, one mating or five matings in a single day. Males were gently aspirated into individual mating vials containing 7ml of 0.75% agar-water. This nutrient-lacking medium was used to remove any potentially confounding influence of a nutritional substrate on mating behaviour, while providing moisture. One virgin female was introduced to each vial for males assigned to the one and five mating treatments and latency to mate and mating duration were recorded by scan sampling approximately every minute. Once mating was complete, the female was removed. For males assigned to the five matings treatment, this process was repeated until five consecutive matings were achieved. If no mating occurred within 60 minutes, the female was replaced with a new virgin female. Matings under five minutes were excluded (n=9) as short matings may not allow complete transfer of sperm and seminal fluids (Gilchrist and Partridge, 2000; Manier *et al*., 2010). Matings over 40 minutes were also excluded (n=4) because they were cases in which males and females had become stuck during sperm transfer (Mason, Rostant and Chapman, 2016). The no mating and one mating treatment vials were set up and handled in the same way as the five mating treatment vials. After the mating assay, experimental males were housed singly in agar-water mating vials overnight.

### Diets and CAFE assay

We used the CAFE assay (Ja et al. 2007) to measure the consumption of protein and carbohydrate liquid synthetic diets (Piper *et al*., 2014; Camus *et al*., 2017, 2018). Recipes are included in Supplementary Tables 1 and 2. Liquid diets included identical volumes of synthetic components (lipids, vitamins, and salts) to create a fully chemically defined diet. The protein diet was complemented with amino acids and the carbohydrate diet was complemented with sucrose. We added a supplement of 20% autoclaved yeast suspension to the protein diet following previous reports that *D. melanogaster* adults reject the pure protein solution (Camus et al., 2017, 2018). The resulting protein diet therefore contained 4% carbohydrate due to the sugar content of the killed yeast.

On the day following the mating assay, experimental males were transferred to fresh agar-water vials in groups of three. Preliminary experiments indicated that individual males ingest too little protein to detect through our approach. Coded labelling was used to ensure that the experimental treatments were anonymised to the observer. Vials were each provided with two 5μl microcapillary tubes (Ringcaps™; Hirschmann Instruments™), one containing the liquid protein diet and the other containing the liquid carbohydrate diet, held in place with a foam bung. Both microcapillary tubes were replaced every 24 hours for five days and the loss of liquid diet was measured from each tube at the meniscus using a digital calliper. We calculated total food consumption as the length of the vector from the origin to the intake values of protein and carbohydrate, in a protein and carbohydrate nutrient space (Supplementary Figure 1) (Camus *et al*., 2018). We calculated relative amounts of protein to carbohydrate ingested as the angle between this vector and the protein axis to give a value between 1-90°. Values of over 45° signify a greater proportion of carbohydrate consumed and values of less than 45° signify a greater proportion of protein consumed. During the experiment, vials were housed in a sealed 50 L lidded box containing a saturated salt solution to create a high humidity environment (77% on average) (Greenspan, 1977) to limit evaporation from the microcapillaries. Ten vials with protein- and carbohydrate-containing microcapillary tubes but without flies were interspersed amidst the experimental vials for each day of the measuring period to track evaporative loss. Mean evaporation was calculated from these vials for both protein and carbohydrate and subtracted from the feeding measures for each day of the measurement period. Instances where evaporation was greater than the measured diet consumption were excluded (n=18) because calculations of dietary preference (angle, see Supplementary Figure 1) require positive intake values. Vials where one or more males escaped or died during the assay were also excluded.

### Reproductive output

All females that mated with experimental males were retained in individual SYA vials seeded with live yeast granules and allowed to lay eggs for approximately four weeks to allow us to profile offspring production over time from each mating. Females were moved to fresh vials every three to five days to ensure larvae would not be food limited. Vials were frozen 13 days after egg-laying to allow all offspring to eclose. The number of adult offspring was counted.

### Ethical note

This research was conducted on fruit flies, which are not subject to any ethical restrictions in the United Kingdom.

### Statistical analysis

Statistical analyses were carried out in R version 4.0.4 (The R Foundation for Statistical Computing, Vienna, Austria, http://www.r-project.org) and all statistical models can be found in Supplementary Table 3.

### Dietary preference and mating duration

We excluded outliers for diet consumption identified by Z-Score calculations (n=15 protein values, n=8 carbohydrate values). Outliers included tubes that had drained of liquid due to accidental contact with a substrate. To account for the effect of block on dietary and mating duration data, we first fitted generalised linear models with block assigned as a fixed factor for each of the response variables length of vector, angle, and duration of mating. Residuals from these models were analysed as response variables in generalised linear mixed models (GLMM) using the R package glmmTMB (Brooks *et al*., 2017) in which treatment and day were included as fixed effects for analysing length of vector and angle, and mate number for mating duration. Individual male id was included as a random effect. We also analysed differences between the treatment groups on day one only to untangle whether there was an effect of the mating treatment on feeding behaviour directly following the mating assay. Data from each block was additionally analysed separately for duration following tests for model fit. Additionally, we included angle and length of vector as response variables in a multivariate analysis of variance (MANOVA) to investigate the effects of treatment and day on the joint response of both variables (Pillai’s trace). Angle and length of vector were centred around a mean of 0 for the MANOVA. Throughout, model fit was checked using the R package DHARMa (Hartig, 2022). Post-hoc pairwise comparisons were carried out on estimated marginal means using the R package emmeans (Lenth *et al*., 2023).

### Reproductive output

To investigate male sperm depletion with multiple mating, we tested the effect of a female’s mate number (i.e., whether the female was the first, second, third, fourth or fifth mate of their male partner) on her offspring output over time (e.g., per vial). For each mated female, we calculated the slope of her adult offspring production regressed against time (i.e., each of the seven sequential 24-hour periods (vials)). Females without data for all seven timepoints were excluded, such as those that died during the data collection period (n=74), as were females that produced no offspring, to exclude reproductive failure events. To account for effects of block, we first fitted a generalised linear model with block as a fixed factor and with individual slopes as the response variable. Residuals from the initial model were entered as the response variable in a linear model against mate number (whether the female was the first, second, third, fourth or fifth mate of their male partner).

### Latency to mate

Latency to mate was analysed as a function of female mate number using the R package survival (Therneau and Grambsch, 2000; Therneau, 2023) and visualised using a Kapalan-Meir curve. Instances where the female partner was replaced were treated as censored values.

### Repeatability analysis

Repeatability of male mating behaviour was analysed for both mating duration and latency for males that mated multiply (three- and five-times mating treatments), by using the R package rptR (Stoffel, Nakagawa and Schielzeth, 2019). Mate number was included as a fixed effect in both models to perform an analysis of enhanced agreement repeatability of male id. Instances of where the female partner was replaced were treated as censored values for repeatability of latency behaviour.

## RESULTS

### Multiple matings do not alter food intake or preference for protein and carbohydrate

The combined total of protein and carbohydrate diet eaten by males (i.e., length of vector) was not significantly different between the 0, 1 or 5 mating treatment groups (Figure 1a) (*χ^2^_2_*=2.12, *P=*0.35). There was no significant interaction between treatment and day (*χ^2^_8_*=5.47, *P=*0.71), suggesting little change in consumption over time. It is possible that an effect of mating on food intake might be strongest immediately after mating. However, the quantity of food consumed on day one was not significantly different among mating treatments (*χ^2^_2_*=1.97, *P=*0.37). There was a significant effect of day on total food consumption (*χ^2^_4_*=11.89, *P=*<0.05), but no consistent effect of day between blocks.

**Figure 1.**
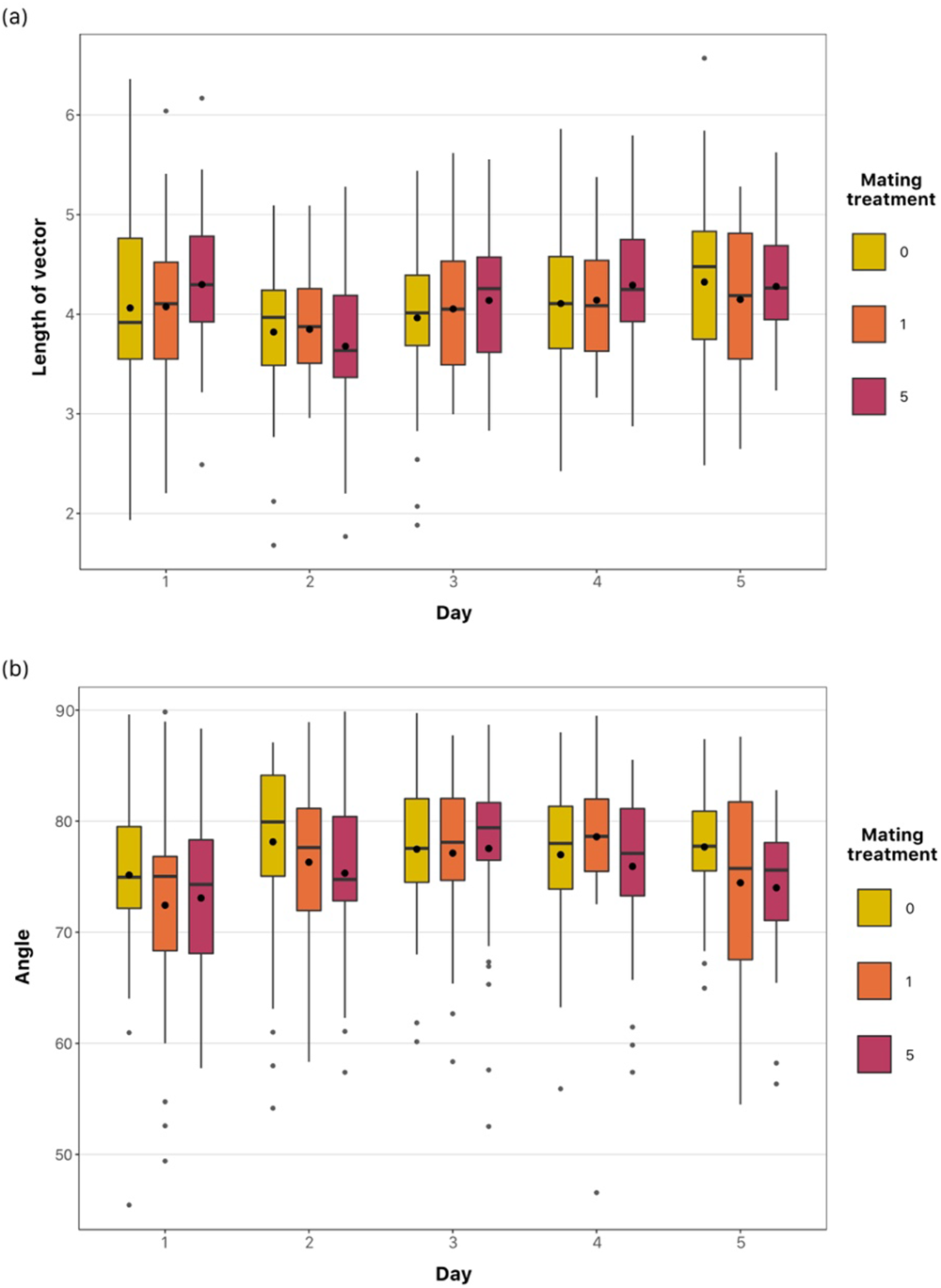
The quantity (a) and composition (b) of food eaten over 24h periods by experimental males assigned to mate 0, 1 or 5 times. (a) Combined consumption of protein and carbohydrate represented by the length of vector, calculated as the distance of an intake value from the origin in P:C nutrient space (see Supplementary Figure 1). (b) Relative composition of protein to carbohydrate eaten (μl), represented by the angle between the length of vector and the protein axis to give a value between 0 – 90°. Angles under 45° indicate that more protein than carbohydrate was consumed, whereas angles over 45° indicate that more carbohydrate than protein was consumed. Boxes represent interquartile range (IQR), with medians as thick horizontal lines and means as large, filled circles. Whiskers represent 1.5 x IQR and small circles represent outliers.

The P:C diet composition (angle) that males ate was not significantly different between 0, 1- or 5-times mated males (*χ^2^_2_*=3.18, *P=*0.20). There was no discernible effect of day (*χ^2^_4_*=4.83, *P=*0.30) and no significant interaction between treatment and day (*χ^2^_8_*=8.7, *P=*0.37) (Figure 1b). Males showed a consistent preference for carbohydrate over protein: all recorded P:C angles were over >45°, indicating a skew to carbohydrate (Supplementary Figure 1). We found no evidence for a stronger effect of mating on diet composition preference immediately after mating (no effect of treatment on day 1 diet composition; *χ^2^_2_*=2.56, *P=*0.28). There was no treatment effect in the raw consumption data for either protein or carbohydrate (Supplementary Figure 2).

We tested whether treatment affected the joint feeding response of both total diet intake and diet composition using MANOVA. This showed no significant effect of treatment (Pillai test statistic *df*_2_=0.0096, approx. *F*_4,1084_=1.31, *P*=0.26) or interaction between treatment and day (Pillai test statistic *df*_8_=0.0219, approx. *F_16, 1084_*=0.75, *P=*0.74) in the multivariate analysis. The joint feeding response varied among days (Pillai test statistic *df*_4_=0.0302, approx. *F*_8,1084_=2.07, *P=*<0.05). These results support the finding that there was no change in the nutrient intake for males as a function of their mating frequency.

### Males become ejaculate-limited after multiple matings

As expected, the total number of offspring sired by a male increased when males mated more than once (*F_2,136_*=49.44, *P=*<0.001) (Figure 2). Males that mated once produced 313 ±SE 16 (*N*=47) adult offspring on average, while an average of 642 ±SE 29 (*N*= 46) and 681 ±SE 38 (*N*=46) offspring were produced by those that mated three and five times, respectively. The number of offspring produced by a male was significantly higher in males that mated three times, or five times compared to those that mated only once (post-hoc Tukey tests: one versus three matings: *t_136_*=-8.08, *P=*<0.001, one versus five matings: *t_136_*=-9.04, *P=*<0.001). However, reproductive output did not differ between the multiply mated groups (*t_136_*=-0.96, *P=*0.61), despite an additional two matings with virgin females. This effect was echoed in an analysis of the slope of decline in the numbers of offspring produced by each female (Figure 3). The slope of decline in female offspring production depended on a female’s position in the mating sequence (*χ^2^_4_*=174.47, *P=*<0.001). Post-hoc Tukey tests showed that this slope of decline was significantly different when comparing across all mate numbers (*P=*<0.01 to *P=<*0.001), with the exception of the fourth versus fifth females to mate with a male (*t_290_*=-1.3, *P=* 0.68). Mating latency and duration were not correlated with offspring production (Supplementary Figure 3).

**Figure 2.**
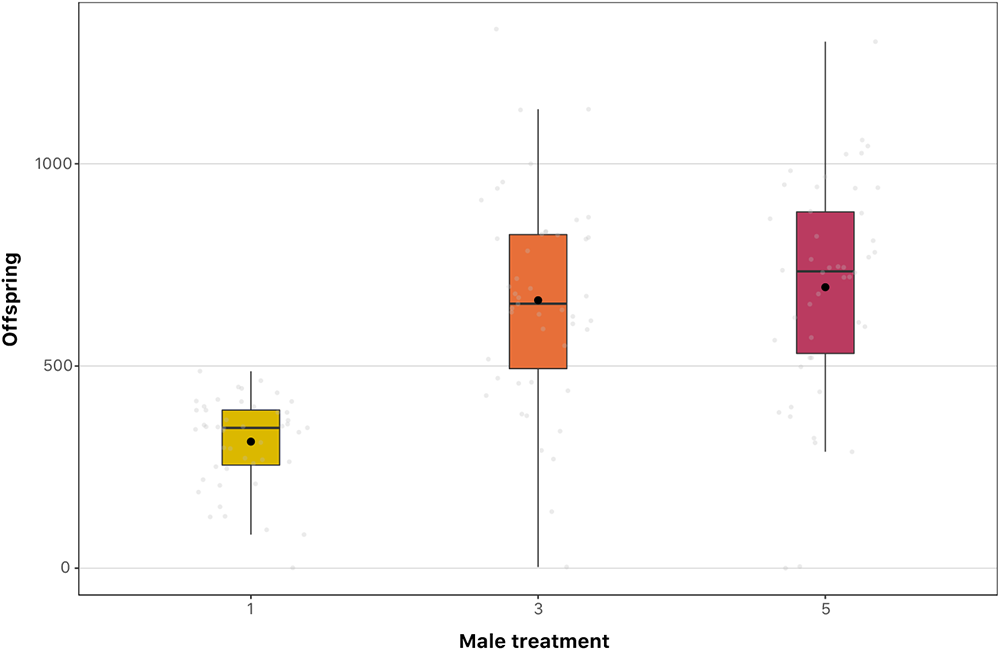
Total numbers of adult offspring sired by males when mated once, three or five times sequentially. Data shown are from experimental blocks one and two where an additional three times mated treatment was included. Boxes represent interquartile range (IQR), with medians as thick horizontal lines and means as large, filled circles. Whiskers represent 1.5 x IQR and small circles are raw data points overlaid on their respective box.

**Figure 3.**
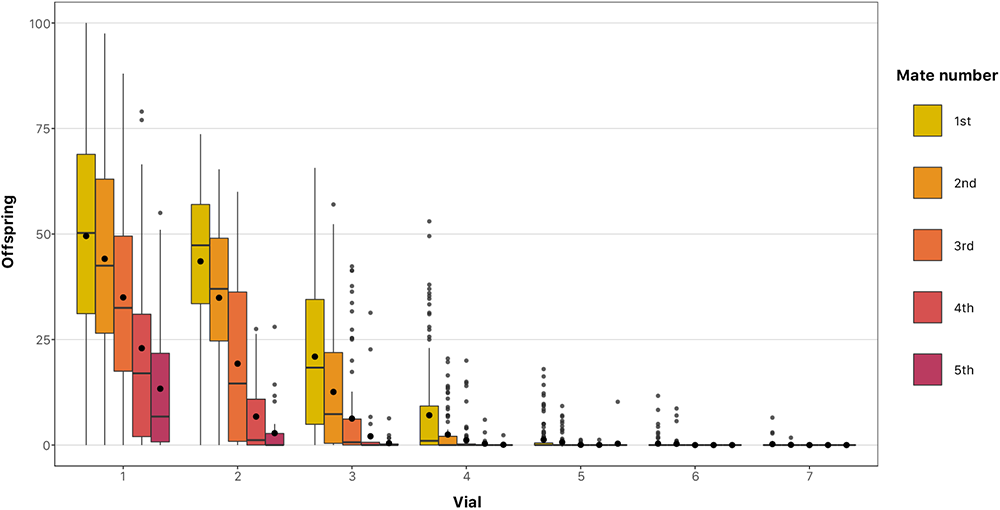
Total numbers of adult offspring produced over 24h by individual females that were a male’s first, second, third, fourth or fifth partner. All females were initially virgins and mated with the experimental treatment males. Measures were taken from vials where females were allowed to lay eggs over three to five days. These periods were normalised to 24h by dividing values by the number of days the female laid eggs in that vial. Boxes represent interquartile range, with medians as thick horizontal lines and means as large, filled circles. Whiskers represent 1.5 x interquartile range and small circles are outliers. Data shown are from blocks one and two where a three times mated treatment was included.

### Males took longer to mate if they had previously mated at least twice

Latency to mate was impacted by how many times a male had previously mated (*χ^2^_4_=*108.65, *P=*<0.001) (Figure 4). The median time to mate was 8.0, 8.5, 11.0, 13.0 and 30.0 minutes for the first, second, third fourth and fifth matings, respectively. Latency between consecutive matings was not significantly different, with the exception of the second versus third mating (post-hoc Tukey: *Z*=3.5, *P=*<0.01), while latency between non-consecutive matings were significantly different from one another (post-hoc Tukey*: P=*<0.001 to *P=*<0.01). There was significant variation in latency between experimental blocks (*χ^2^_3_=*21.31, *P=*<0.001). Males also showed an increased reluctance to mate later in the mating sequence, as evident in the requirement to add additional virgin females for mating to occur (Supplementary Figure 4). Latency was not a repeatable individual behaviour (repeatability score; *R*=0.02, *P=*0.19).

**Figure 4.**
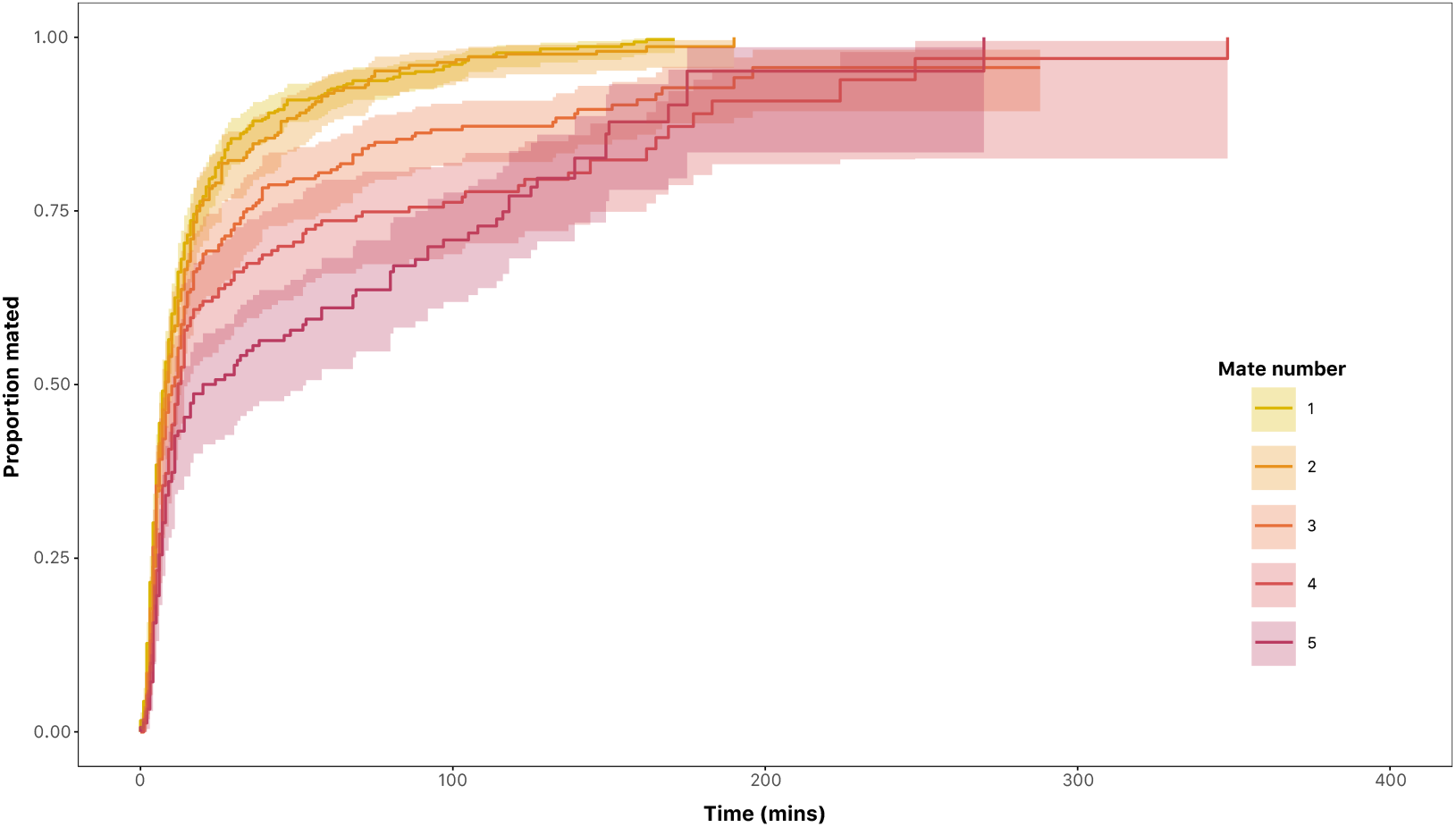
Kaplan-Meier survival curves illustrating the proportion of experimental males initiating their first, second, third, fourth or fifth matings. Solid lines (with surrounding confidence intervals in matching colourways) represent each of the one to five sequential mating treatments, in which males were presented with a new virgin female. Data shown are taken from all four blocks.

### Mating duration decreased after the first mating

Mating duration was significantly altered by mate number (*χ^2^*_4_=190.39, *P=*<0.001) (Figure 5) in a model run across data from all blocks. Significant deviation from the expected distribution of residuals was detected during model fit tests, therefore blocks were also analysed separately: mate number had a significant effect on duration in all blocks (*P=*<0.001 in all cases). Pairwise comparisons showed first matings were significantly longer than subsequent matings (post-hoc Tukey tests: *P=*<0.001 to <0.05) with the exception of the fourth mating in both block 1 (*t_168_*=2.30, *P=*0.149) and block 2 (*t_178_*=2.60, *P=*0.075). All matings after the first did not differ from each other. There was significant but minimal repeatability in mating duration across the multiple matings of an individual male, once accounting for the variation caused by mate number (repeatability score; *R*=0.152, *P=*<0.001).

**Figure 5.**
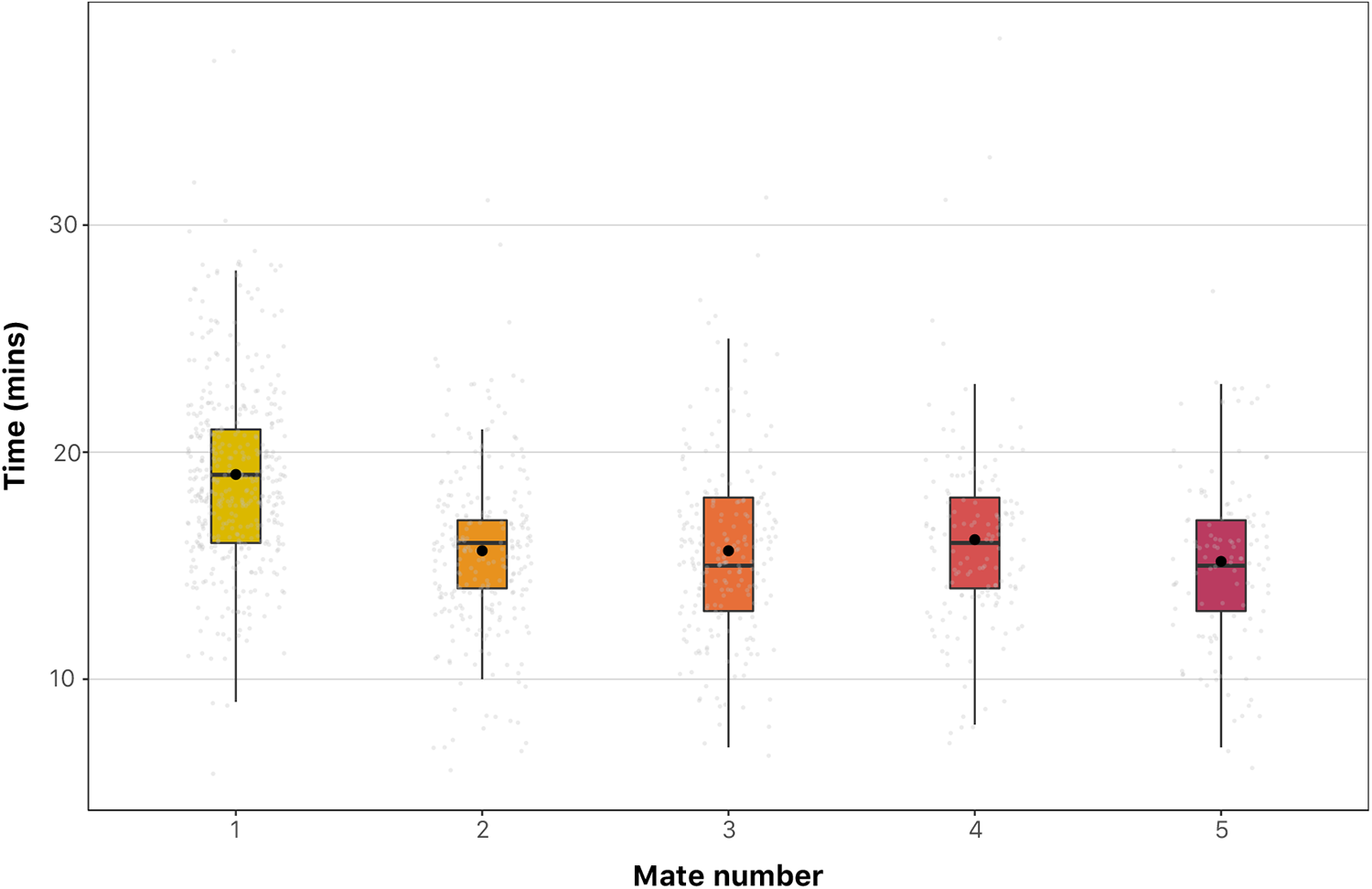
Mating duration of individual males when mating for the first, second, third, fourth or fifth time with a virgin female in a single day. Boxes represent interquartile range (IQR), with median shown as the thick horizontal line and mean shown as the large, filled circle. Whiskers represent 1.5 x IQR and jittered raw data are overlaid on each box. Data shown are taken from all blocks.

## DISCUSSION

Though it is often assumed that production of sperm incurs little cost to males, ejaculate production costs as a whole can be significant (Dewsbury, 1982). Nutritional demands due to sperm depletion may not be discernible after a single mating (Perry and Tse, 2013; Camus *et al*., 2018). However, they are predicted to be evident after multiple matings, which males of many species experience in natural settings. To our knowledge, this broad prediction had not previously been tested in species lacking nuptial gifts. We tested whether mating multiply, would trigger male fruit flies to alter their dietary preference to replenish ejaculate components. We tested 3 predictions. Consistent with prediction 1, we found that males in the high mating rate treatment were more ejaculate depleted. However, predictions 2 and 3 were not supported: we did not detect evidence for a change in macronutrient preference, nor did males eat more, in the high mating rate treatment compared to the unmated and low mating rate treatments.

### Male ejaculate depletion with consecutive matings

We found that males were severely depleted in their ability to transfer ejaculates with fertilisation potential to females by their fifth mating. This was observed in the pattern of offspring production of their mates, with females that mated later in a male’s mating sequence producing no or markedly reduced numbers of adult offspring compared with the first mate. Furthermore, the reduction in offspring produced showed a non-linear mate number-dependant decline. This is in accord with results on reproductive success in multiply mated males observed previously (Hihara, 1981; Linklater *et al*., 2007; Douglas, Anderson and Saltz, 2020; Macartney *et al*., 2021) . This effect is likely to result chiefly from seminal fluid rather than sperm depletion because it has been observed that males retain some sperm in their seminal vesicles, but lack fluid in the accessory gland, after four to five matings (Lefevre and Jonsson, 1962; Gillott, 2003; Macartney *et al*., 2021; see also Reinhardt, Naylor and Siva-Jothy, 2011). The case is similar in the common bedbug, *Cimex lectularius*, where reserves of seminal fluid declined faster than sperm reserves in the male’s reproductive organs and seminal fluid depletion limited male remating (Reinhardt, Naylor and Siva-Jothy, 2011). An increased reluctance to mate was also evident in males as they mated in sequence: males took longer to mating after the first two matings, later matings showed an increase in male refusal to mate, and fifth matings required more instances of swapping in new virgin females to get matings to occur. Duration of mating was also significantly shorter in all matings after a male’s first, suggesting a reduced investment in subsequent matings.

### Ejaculate depletion does not alter male macronutrient intake or preference

We predicted that following the depletion of ejaculate reserves after five matings, males would increase consumption of food, and especially protein, to allow for more efficient regeneration of proteinaceous sperm and seminal peptides (Perry, Sirot and Wigby, 2013). Yeast availability in adult male diet, as a protein supplement, has been found to influence a male’s ability to gain mates and sire offspring (Fricke, Bretman and Chapman, 2008). Despite the importance of protein intake, males were either unable to translate their physiological requirements into dietary preference or gained no benefit from doing so, and hence males remained on similar P:C trajectories with similar food intake rates regardless of their mating rate. Dietary preference was consistent between singly mated and unmated male *D. melanogaster,* which is congruent with both previous work (Camus *et al*., 2018) and with the observation that levels of seminal fluid are not significantly depleted after a single mating (Lefevre and Jonsson, 1962).

Males in this study favoured a low P:C in their diet in all mating treatments, at a ratio of 1:4. This is consistent with previous studies of male *D. melanogaster* (Ribeiro and Dickson, 2010; Lee, Kim and Min, 2013; Jensen *et al*., 2015; Camus *et al*., 2018) and other male insects (e.g., field crickets, *Teleogryllus oceanicus* and German cockroaches, *B. germanica*) (Jensen and Silverman, 2018; Ng, Simpson and Simmons, 2019). Though both sexes of *D. melanogaster* prefer a carbohydrate-biased diet, male fruit flies exhibit an even greater preference for carbohydrate than females do. This is thought to result from the demands of performing energetically costly courtship rituals, which are essential for male reproductive success (Bastock and Manning, 1955; von Schilcher, 1976; Lee, Kim and Min, 2013; Camus *et al*., 2018). When placed with non-receptive females, males continue courting for hours (Bastock and Manning, 1955). It is therefore possible that males might increase carbohydrate intake after multiple matings, to recoup nutritional resources lost to extra bouts of courtship. However, this prediction was not upheld, as we observed that carbohydrate intake was unchanged across the treatment groups.

The results from this study suggest that male *D. melanogaster* choose a ratio of P:C and intake rate that is insensitive to their sexual experience. Some invariance in male diet preference is also suggested by a previous study of dietary preference following macronutrient deprivation (Ribeiro and Dickson, 2010). After maintenance on a sucrose-only diet, females strongly preferred yeast after only three days on sucrose, while males took 10 days to reach an equivalent yeast preference and lost this preference far more rapidly when returned to a yeast medium (Ribeiro and Dickson, 2010). That study provides some evidence that males are able to respond to a severe protein deficit but suggests that males do so more slowly than do females. The absence of evidence in our study for increased food or protein ingestion by multiply mated males suggests that multiple matings do not induce such extreme protein limitation in males, even near the physiological limits for mating rate. In another study, five-times mated male *D. melanogaster* sire a number of offspring equivalent to their first mating, provided that the fifth mating took place after a 24h respite period (Loyau *et al*., 2012). However, males in this study remained on a solid cornmeal-agar-yeast diet, so it is unclear whether males altered their food intake over the 24h respite period. Another study investigated the reproductive output of multiply mated males over 4h mating bouts on successive days (Douglas, Anderson and Saltz, 2020). Males were able to remate multiply on consecutive days but did not necessarily sire more offspring, suggesting incomplete regeneration of ejaculate reserves between days (Douglas, Anderson and Saltz, 2020). The nutritional intake patterns of males that sustain a high mating rate over multiple days would be interesting to investigate further.

Sexual selection theory suggests that males typically maximise reproductive success by mating as many times as possible (Andersson, 1994; Arnqvist and Rowe, 2005). However, in natural settings, multiple factors may affect a male’s ability to gain mating and remating opportunities, including access to females, female mate preferences, and male-male competition. It may be that males have not evolved a mechanism to cope with sudden ejaculate depletion because the opportunity for consecutive, multiple rematings is rare in nature. There is also evidence that a mated male is less attractive as a mate to females (Loyau *et al*., 2012; Douglas, Anderson and Saltz, 2020). Hence, in scenarios where access to females is unlimited, males may gain fitness by mating at maximal frequencies regardless of ejaculate depletion. Consistent with this idea, we observed that both multiply mated treatment groups had higher offspring numbers than those mated only mated once, while five times mated males were still able to sire offspring with their last mate and had a marginally higher reproductive output than three-times mated males (though not significantly higher). More data on male *D. melanogaster* remating behaviour in natural contexts would be helpful for exploring these ideas further.

Our results contrast markedly with findings in females, which respond to nutritional deficit and the initiation of reproduction and dynamically adjust their intake accordingly. Specifically, *D. melanogaster* females alter their diet preference from a low P:C diet as virgins (from a level that is similar to the observed P:C ratio of males) to a higher P:C ratio after mating (Barnes *et al*., 2008; Ribeiro and Dickson, 2010; Lee, Kim and Min, 2013; Corrales-Carvajal, Faisal and Ribeiro, 2016; Camus *et al*., 2018). This sex differences is likely to ultimately result from contrasting reproductive strategies between the sexes, whereby females might gain most reproductive success from limiting their number of mates and have greater nutritional requirements to support offspring production. Proximate explanations for the sex differences in dietary responses to mating include the mediation of female response by the sex peptide protein transferred from males to females during mating (Chapman *et al*., 2003; Carvalho *et al*., 2006; Barnes *et al*., 2008; Ribeiro and Dickson, 2010; Apger-McGlaughon and Wolfner, 2013) and gene expression differences in nutrient sensing pathways between the sexes (Bennett-Keki *et al*., 2023).

In conclusion, we show that multiply mated *D. melanogaster* males do not, or cannot, adjust their intake of protein and carbohydrate diets compared with unmated or singly mated males. Surprisingly, this lack of dietary response occurs even despite significant ejaculate depletion. This study highlights the gap in knowledge regarding male nutrient homeostasis in response to sexual experience, in contrast to the data available on females. Males of many insect species adjust their ejaculate investment in response to the sexual environment (Gage, 1991; Gage and Barnard, 1996). However, our data suggest that males don’t support this shift in ejaculate allocation by responding to any nutrient debt it entails. In addition, since five-times-mated male *D. melanogaster* might still have motile sperm present in their vesicles (Lefevre and Jonsson, 1962; Gillott, 2003), perhaps *D. melanogaster* do not perceive a protein requirement until sperm reserves have been fully depleted. Overall, the results suggest that males remain on a fixed feeding trajectory even when mating close to their daily functional maxima, and do not increase nutrient intake to recoup their reduced ability to transfer ejaculates to females.

## DATA ACCESSIBILITY

Data are provided in additional supplementary information for review.

## AUTHOR CONTRIBUTIONS

M.C.S. and J.C.P. conceived and designed the study; M.C.S. conducted the experiments with assistance from J.C.P.; M.C.S. performed data analysis and visualisation with advice and supervision from T.C. and J.C.P.; all authors wrote the original draft, and edited and approved the final draft for publication.

## COMPETING INTERESTS

The authors declare no conflicts of interest.

## FUNDING

This work was supported by a UK NERC PhD studentship held by M.C.S. within the ARIES Doctoral Training Partnership (NE/S007334/1), NERC grant (NE/P017193/1) held by J.C.P., and NERC grant (NE/R000891/1) and BBSRC grant (BB/W005174/1) held by T.C.

## ACKNOWLEDGMENTS

We thank Wayne Rostant for his advice on statistical modelling, help with a mating assay and general discussion in the early evaluations of this study. We also thank Lauren Harrison for further support on model fitting, Ginny Greenway for advice on repeatability analysis and helpful comments on the study, and Paul Candon and Kerri Armstrong for technical assistance.

## Notes

### Competing Interest Statement

The authors have declared no competing interest.

## REFERENCES

Andersson, M. (1994) Sexual Selection. Princeton University Press.

Apger-McGlaughon, J. and Wolfner, M.F. (2013) ‘Post-mating change in excretion by mated Drosophila melanogaster females is a long-term response that depends on sex peptide and sperm’, Journal of Insect Physiology, 59(10), pp. 1024–1030. Available at: https://doi.org/10.1016/j.jinsphys.2013.07.001.

Arnqvist, G. and Rowe, L. (2005) Sexual Conflict. Available at: https://press.princeton.edu/books/paperback/9780691122182/sexual-conflict (Accessed: 23 March 2023).

Barnes, A.I. et al. (2008) ‘Feeding, fecundity and lifespan in female Drosophila melanogaster’, Proceedings of the Royal Society B: Biological Sciences, 275(1643), pp. 1675–1683. Available at: https://doi.org/10.1098/rspb.2008.0139.

Bastock, M. and Manning, A. (1955) ‘The Courtship of Drosophila Melanogaster’, Behaviour, 8(1), pp. 85–110. Available at: https://doi.org/10.1163/156853955X00184.

Bateman, A.J. (1948) ‘Intra-sexual selection in Drosophila’, Heredity, 2(3), pp. 349–368. Available at: https://doi.org/10.1038/hdy.1948.21.

Bennett-Keki, S. et al. (2023) ‘Sex-biased gene expression in nutrient-sensing pathways’, Proceedings of the Royal Society B: Biological Sciences, 290(1994), p. 20222086. Available at: https://doi.org/10.1098/rspb.2022.2086.

Bownes, M. and Blair, M. (1986) ‘The effects of a sugar diet and hormones on the expression of the Drosophila yolk-protein genes’, Journal of Insect Physiology, 32(5), pp. 493–501. Available at: https://doi.org/10.1016/0022-1910(86)90011-9.

Brooks, M., E., et al. (2017) ‘glmmTMB Balances Speed and Flexibility Among Packages for Zero-inflated Generalized Linear Mixed Modeling’, The R Journal, 9(2), p. 378. Available at: https://doi.org/10.32614/RJ-2017-066.

Camus, M.F. et al. (2017) ‘Sex and genotype effects on nutrient-dependent fitness landscapes in *Drosophila melanogaster*’, Proceedings of the Royal Society B: Biological Sciences, 284(1869), p. 20172237. Available at: https://doi.org/10.1098/rspb.2017.2237.

Camus, M.F. et al. (2018) ‘Dietary choices are influenced by genotype, mating status, and sex in Drosophila melanogaster’, Ecology and Evolution, 8(11), pp. 5385–5393. Available at: https://doi.org/10.1002/ece3.4055.

Carey, M.R. et al. (2022) ‘Mapping sex differences in the effects of protein and carbohydrates on lifespan and reproduction in Drosophila melanogaster: is measuring nutrient intake essential?’, Biogerontology, 23(1), pp. 129–144. Available at: https://doi.org/10.1007/s10522-022-09953-2.

Carvalho, G.B. et al. (2006) ‘Allocrine modulation of feeding behavior by the Sex Peptide of Drosophila’, Current biology: CB, 16(7), pp. 692–696. Available at: https://doi.org/10.1016/j.cub.2006.02.064.

Chapman, T. et al. (2003) ‘The sex peptide of Drosophila melanogaster: Female post-mating responses analyzed by using RNA interference’, Proceedings of the National Academy of Sciences, 100(17), pp. 9923–9928. Available at: https://doi.org/10.1073/pnas.1631635100.

Chapman, T., Trevitt, S. and Partridge, L. (1994) ‘Remating and male-derived nutrients in Drosophila melanogaster’, Journal of Evolutionary Biology, 7(1), pp. 51–69. Available at: https://doi.org/10.1046/j.1420-9101.1994.7010051.x.

Corrales-Carvajal, V.M., Faisal, A.A. and Ribeiro, C. (2016) ‘Internal states drive nutrient homeostasis by modulating exploration-exploitation trade-off’, eLife. Edited by I.D. Couzin, 5, p. e19920. Available at: https://doi.org/10.7554/eLife.19920.

De Crespigny, F.E.C., Pitt, T.D. and Wedell, N. (2006) ‘Increased male mating rate in Drosophila is associated with Wolbachia infection’, Journal of Evolutionary Biology, 19(6), pp. 1964–1972. Available at: https://doi.org/10.1111/j.1420-9101.2006.01143.x.

Dewsbury, D.A. (1982) ‘Ejaculate Cost and Male Choice’, The American Naturalist, 119(5), pp. 601– 610.

Douglas, T., Anderson, R. and Saltz, J.B. (2020) ‘Limits to male reproductive potential across mating bouts in Drosophila melanogaster’, Animal Behaviour, 160, pp. 25–33. Available at: https://doi.org/10.1016/j.anbehav.2019.11.009.

Drummond-Barbosa, D. and Spradling, A.C. (2001) ‘Stem Cells and Their Progeny Respond to Nutritional Changes during Drosophila Oogenesis’, Developmental Biology, 231(1), pp. 265–278. Available at: https://doi.org/10.1006/dbio.2000.0135.

Fanson, B.G. et al. (2009) ‘Nutrients, not caloric restriction, extend lifespan in Queensland fruit flies (Bactrocera tryoni)’, Aging Cell, 8(5), pp. 514–523. Available at: https://doi.org/10.1111/j.1474-9726.2009.00497.x.

Fricke, C., Bretman, A. and Chapman, T. (2008) ‘Adult Male Nutrition and Reproductive Success in Drosophila Melanogaster’, Evolution, 62(12), pp. 3170–3177. Available at: https://doi.org/10.1111/j.1558-5646.2008.00515.x.

Gage, A.R. and Barnard, C.J. (1996) ‘Male crickets increase sperm number in relation to competition and female size’, Behavioral Ecology and Sociobiology, 38(5), pp. 349–353. Available at: https://doi.org/10.1007/s002650050251.

Gage, M.J.G. (1991) ‘Risk of sperm competition directly affects ejaculate size in the Mediterranean fruit fly’, Animal Behaviour, 42(6), pp. 1036–1037. Available at: https://doi.org/10.1016/S0003-3472(05)80162-9.

Gilchrist, A.S. and Partridge, L. (2000) ‘Why It Is Difficult to Model Sperm Displacement in Drosophila Melanogaster: The Relation Between Sperm Transfer and Copulation Duration’, Evolution, 54(2), pp. 534–542. Available at: https://doi.org/10.1111/j.0014-3820.2000.tb00056.x.

Gillott, C. (2003) ‘Male Accessory Gland Secretions: Modulators of Female Reproductive Physiology and Behavior’, Annual Review of Entomology, 48(1), pp. 163–184. Available at: https://doi.org/10.1146/annurev.ento.48.091801.112657.

Greenspan, L. (1977) ‘Humidity Fixed Points of Binary Saturated Aqueous Solutions’, *Journal of Research of the National Bureau of Standards. Section A*, Physics and Chemistry, 81A(1), pp. 89–96. Available at: https://doi.org/10.6028/jres.081A.011.

Hartig, F. (2022) ‘DHARMa: Residual Diagnostics for Hierarchical (Multi-Level / Mixed) Regression Models’. Available at: https://CRAN.R-project.org/package=DHARMa.

Hihara, F. (1981) ‘Effects of the Male Accessory Gland Secretion on Oviposition and Remating in Females of Drosophila melanogaster’, 動物学雑誌 Zoological Magazine, 90(3), pp. 307–316. Available at: https://doi.org/10.34435/zm004883.

Ja, W.W. et al. (2007) ‘Prandiology of Drosophila and the CAFE assay’, Proceedings of the National Academy of Sciences, 104(20), pp. 8253–8256. Available at: https://doi.org/10.1073/pnas.0702726104.

Jensen, K. et al. (2015) ‘Sex-specific effects of protein and carbohydrate intake on reproduction but not lifespan in Drosophila melanogaster’, Aging Cell, 14(4), pp. 605–615. Available at: https://doi.org/10.1111/acel.12333.

Jensen, K. and Silverman, J. (2018) ‘Frequently mated males have higher protein preference in German cockroaches’, Behavioral Ecology, 29(6), pp. 1453–1461. Available at: https://doi.org/10.1093/beheco/ary104.

Lee, K.P. et al. (2008) ‘Lifespan and reproduction in Drosophila: New insights from nutritional geometry’, Proceedings of the National Academy of Sciences of the United States of America, 105(7), pp. 2498–2503. Available at: https://doi.org/10.1073/pnas.0710787105.

Lee, K.P., Kim, J.-S. and Min, K.-J. (2013) ‘Sexual dimorphism in nutrient intake and life span is mediated by mating in Drosophila melanogaster’, Animal Behaviour, 86(5), pp. 987–992. Available at: https://doi.org/10.1016/j.anbehav.2013.08.018.

Lefevre, G. and Jonsson, U.B. (1962) ‘Sperm Transfer, Storage, Displacement, and Utilization in Drosophila Melanogaster’, Genetics, 47(12), pp. 1719–1736.

Lenth, R.V. et al. (2023) ‘emmeans: Estimated Marginal Means, aka Least-Squares Means’. Available at: https://CRAN.R-project.org/package=emmeans (Accessed: 6 March 2023).

Leshner, A.I., Siegel, H.I. and Collier, G. (1972) ‘Dietary self-selection by pregnant and lactating rats’, Physiology & Behavior, 8(1), pp. 151–154. Available at: https://doi.org/10.1016/0031-9384(72)90144-8.

Linklater, J.R. et al. (2007) ‘EJACULATE DEPLETION PATTERNS EVOLVE IN RESPONSE TO EXPERIMENTAL MANIPULATION OF SEX RATIO IN DROSOPHILA MELANOGASTER’, Evolution, 61(8), pp. 2027–2034. Available at: https://doi.org/10.1111/j.1558-5646.2007.00157.x.

Loyau, A. et al. (2012) ‘When not to copy: female fruit flies use sophisticated public information to avoid mated males’, Scientific Reports, 2(1), p. 768. Available at: https://doi.org/10.1038/srep00768.

Macartney, E.L. et al. (2021) ‘Sperm depletion in relation to developmental nutrition and genotype in Drosophila melanogaster’, Evolution, 75(11), pp. 2830–2841. Available at: https://doi.org/10.1111/evo.14373.

Maklakov, A.A. et al. (2008) ‘Sex-Specific Fitness Effects of Nutrient Intake on Reproduction and Lifespan’, Current Biology, 18(14), pp. 1062–1066. Available at: https://doi.org/10.1016/j.cub.2008.06.059.

Manier, M.K. et al. (2010) ‘Resolving Mechanisms of Competitive Fertilization Success in Drosophila melanogaster’, Science, 328(5976), pp. 354–357. Available at: https://doi.org/10.1126/science.1187096.

Mason, J.S., Rostant, W.G. and Chapman, T. (2016) ‘Resource limitation and responses to rivals in males of the fruit fly Drosophila melanogaster’, Journal of Evolutionary Biology, 29(10), pp. 2010– 2021. Available at: https://doi.org/10.1111/jeb.12924.

Newell, N.R. et al. (2020) ‘The Drosophila Post-mating Response: Gene Expression and Behavioral Changes Reveal Perdurance and Variation in Cross-Tissue Interactions’, G3: Genes |Genomes|Genetics, 10(3), pp. 967–983. Available at: https://doi.org/10.1534/g3.119.400963.

Ng, S.H., Simpson, S.J. and Simmons, L.W. (2019) ‘Sex differences in nutrient intake can reduce the potential for sexual conflict over fitness maximization by female and male crickets’, Journal of Evolutionary Biology, 32(10), pp. 1106–1116. Available at: https://doi.org/10.1111/jeb.13513.

Olsson, M., Madsen, T. and Shine, R. (1997) ‘Is sperm really so cheap? Costs of reproduction in male adders, Vipera berus’, Proceedings of the Royal Society of London. Series B: Biological Sciences, 264(1380), pp. 455–459. Available at: https://doi.org/10.1098/rspb.1997.0065.

Perry, J.C. (2011) ‘Mating stimulates female feeding: testing the implications for the evolution of nuptial gifts’, Journal of Evolutionary Biology, 24(8), pp. 1727–1736. Available at: https://doi.org/10.1111/j.1420-9101.2011.02299.x.

Perry, J.C., Sirot, L. and Wigby, S. (2013) ‘The seminal symphony: how to compose an ejaculate’, Trends in Ecology & Evolution, 28(7), pp. 414–422. Available at: https://doi.org/10.1016/j.tree.2013.03.005.

Perry, J.C. and Tse, C.T. (2013) ‘Extreme Costs of Mating for Male Two-Spot Ladybird Beetles’, PLOS ONE, 8(12), p. e81934. Available at: https://doi.org/10.1371/journal.pone.0081934.

Piper, M.D.W. et al. (2014) ‘A holidic medium for Drosophila melanogaster’, Nature Methods, 11(1), pp. 100–105. Available at: https://doi.org/10.1038/nmeth.2731.

Reddiex, A.J. et al. (2013) ‘Sex-Specific Fitness Consequences of Nutrient Intake and the Evolvability of Diet Preferences.’, The American Naturalist, 182(1), pp. 91–102. Available at: https://doi.org/10.1086/670649.

Reinhardt, K., Naylor, R. and Siva-Jothy, M.T. (2011) ‘Male Mating Rate Is Constrained by Seminal Fluid Availability in Bedbugs, Cimex lectularius’, PLOS ONE, 6(7), p. e22082. Available at: https://doi.org/10.1371/journal.pone.0022082.

Ribeiro, C. and Dickson, B.J. (2010) ‘Sex Peptide Receptor and Neuronal TOR/S6K Signaling Modulate Nutrient Balancing in Drosophila’, Current Biology, 20(11), pp. 1000–1005. Available at: https://doi.org/10.1016/j.cub.2010.03.061.

Richter, C., P. and Barelare, B., JR. (1938) ‘Nutritional requirements of pregnant and lactating rats studied by the self selection method’, Endocrinology, 23(1), pp. 15–24. Available at: https://doi.org/10.1210/endo-23-1-15.

von Schilcher, F. (1976) ‘The role of auditory stimuli in the courtship of Drosophila melanogaster’, Animal Behaviour, 24(1), pp. 18–26. Available at: https://doi.org/10.1016/S0003-3472(76)80095-4.

Simpson, S.J., Le Couteur, D.G. and Raubenheimer, D. (2015) ‘Putting the Balance Back in Diet’, Cell, 161(1), pp. 18–23. Available at: https://doi.org/10.1016/j.cell.2015.02.033.

Simpson, S.J. and Raubenheimer, D. (1997) ‘Geometric Analysis of Macronutrient Selection in the Rat’, Appetite, 28(3), pp. 201–213. Available at: https://doi.org/10.1006/appe.1996.0077.

Simpson, S.J. and Raubenheimer, D. (2012) The Nature of Nutrition: A Unifying Framework from Animal Adaptation to Human Obesity. Princeton University Press.

Simpson, S.J., Ribeiro, C. and González-Tokman, D. (2018) ‘Feeding behavior’, in A. Córdoba-Aguilar, D. González-Tokman, and I. González-Santoyo (eds) Insect Behavior: From Mechanisms to Ecological and Evolutionary Consequences. Oxford University Press, p. 0. Available at: https://doi.org/10.1093/oso/9780198797500.003.0008.

Stoffel, M.A., Nakagawa, S. and Schielzeth, H. (2019) ‘An introduction to repeatability estimation with rptR’. Available at: https://cran.r-project.org/web/packages/rptR/vignettes/rptR.html (Accessed: 16 March 2023).

Therneau, T.M. (2023) ‘A Package for Survival Analysis in R’. Available at: https://CRAN.R-project.org/package=survival (Accessed: 6 March 2023).

Therneau, T.M. and Grambsch, P.M. (2000) Modeling Survival Data: Extending the Cox Model. New York: Springer.

Trivers, R.L. (1972) ‘Parental Investment and Sexual Selection’, in B. Campbell (ed.) Sexual Selection and the Descent of Man: The Darwinian Pivot. Somerset, UNITED STATES: Taylor & Francis Group, p. 137. Available at: http://ebookcentral.proquest.com/lib/uea/detail.action?docID=4926119 (Accessed: 26 May 2022).

