## Supplementary Information for "Nutritional insensitivity to mating in male fruit flies"

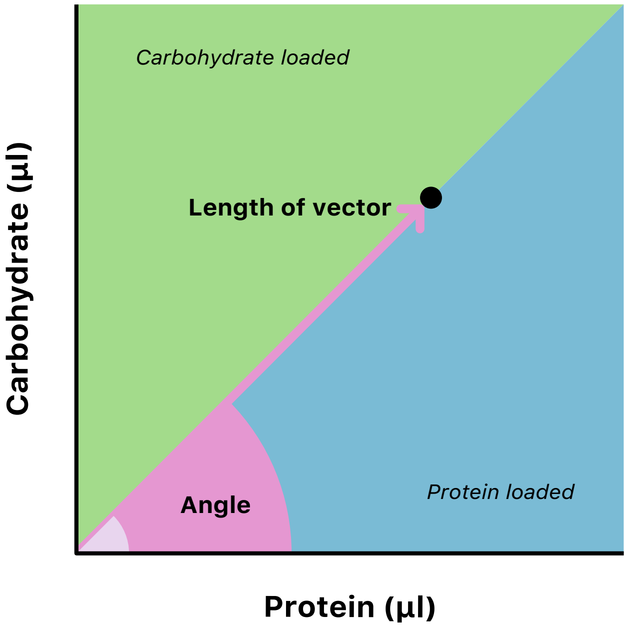


Supplementary Figure 1

Graphical representation of the trigonometric conversion used to calculate angle and length of vector from raw feeding data. Protein and carbohydrate intake is represented by the large black circle. The length of vector for each intake is calculated using distance from the origin (0,0). Points further from the origin signify greater total consumption of macronutrients. The angle between the vector and the x-axis represents composition. Angle values less than 45° denote a greater proportion of protein (the blue space) and values greater than 45° denote a greater proportion of carbohydrate (the green space) eaten.

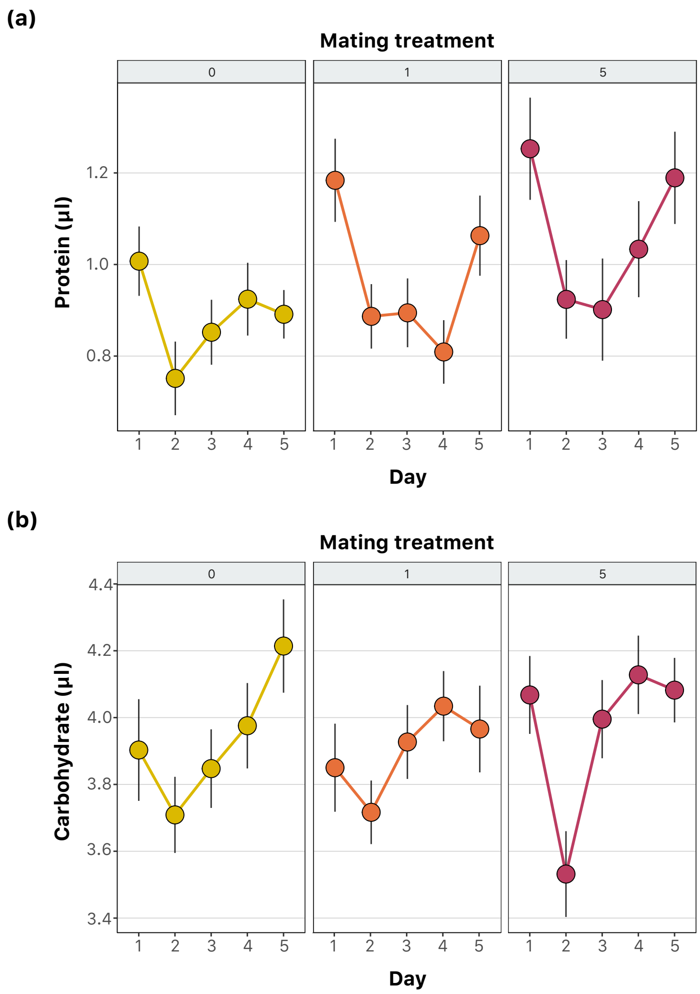


Supplementary Figure 2

Mean (±SE) intake (µl) of (a) protein and (b) carbohydrate eaten over 24h periods by experimental males mated 0, 1 or 5 times. Experimental males were kept in vials in groups of three and presented with a choice of protein and carbohydrate synthetic liquid diets. Black vertical lines denote standard error around the mean (large, filled circles). (a) Raw protein intake was not significantly affected by treatment, day, or the interaction between treatment and day (treatment: $\chi^2_2$= 4.93, *P*=0.08; day: $\chi^2_4$= 6.25, *P*=0.18; treatment*day: $\chi^2_8$= 8.22, *P*=0.41). Treatment was not significant on day 1 only ($\chi^2_2$= 4.17, *P*=0.12). (b) Raw carbohydrate intake was not significantly affected by treatment or the interaction between treatment and day (treatment: $\chi^2_2$= 1.6, *P*=0.45; treatment*day: $\chi^2_8$=6.42, *P*=0.60). There was a significant effect of day ($\chi^2_4$=11.78, *P*=<0.05). Treatment was not significant on day 1 only ($\chi^2_2$= 1.35, *P*=0.51).

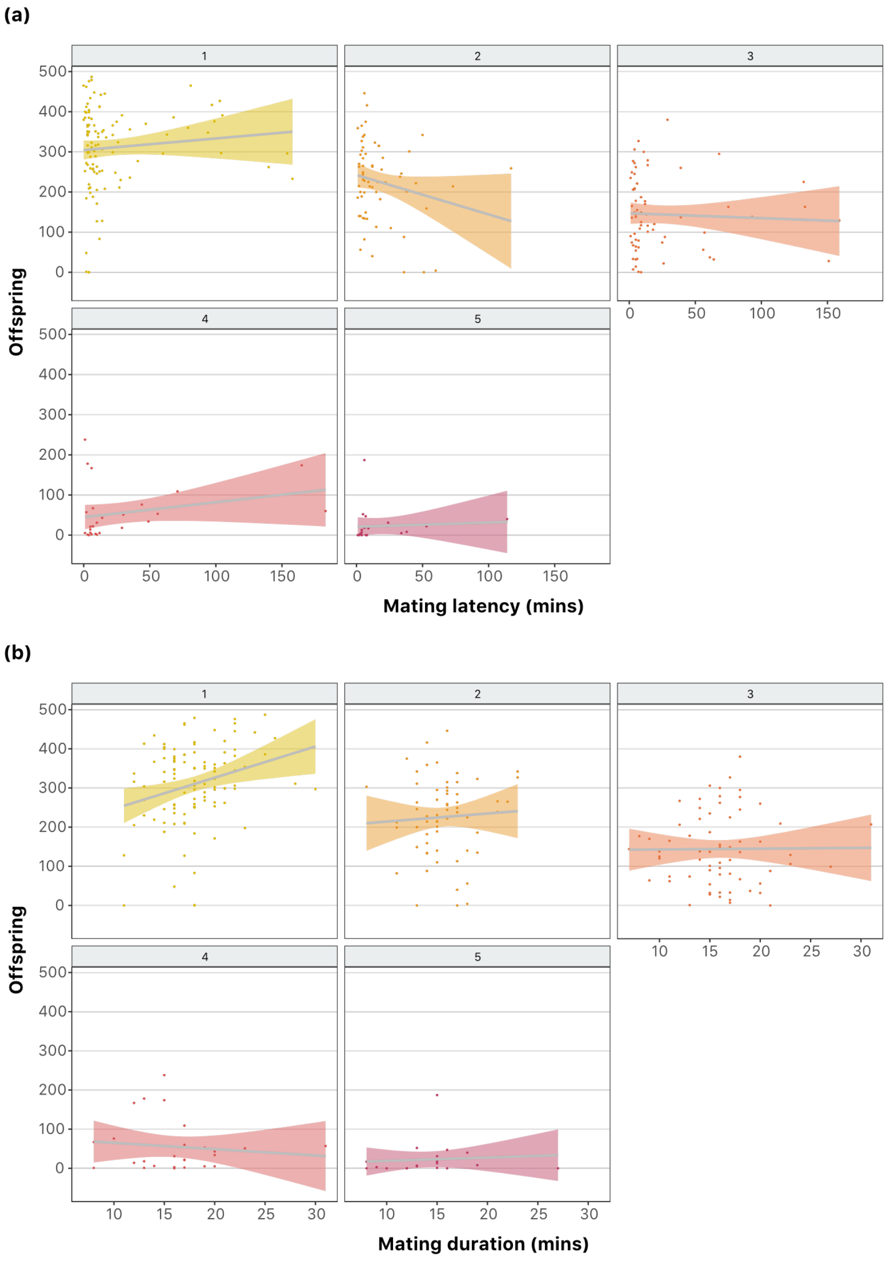


Supplementary Figure 3

Relationship between mating traits and the offspring produced from a single mating. Data shows the offspring produced from a single mating between a male and female fly, where females were mated as virgins and were the first, second, third, fourth or fifth female to mate with a male partner, against the (a) latency to mate and (b) duration of each mating. Raw data points are shown as small circles and linear regression lines are overlaid in grey with coloured confidence intervals. Statistical analysis was carried out with transformed latency and duration data centred around a mean of 0. Preliminary modelling showed an insignificant effect of block, which was subsequently removed from the model. There was no significant interaction between duration, latency and mate number when included as a three-way interaction in a generalised linear mixed model (duration*latency*mate number; $p$=0.41). Model testing showed a reduced model to have the best fit, with mate number having a significant effect on offspring production ($\chi^2_4$=281.77, $p$=<0.001), but not duration ($p$=0.07) or latency ($p$=0.96).

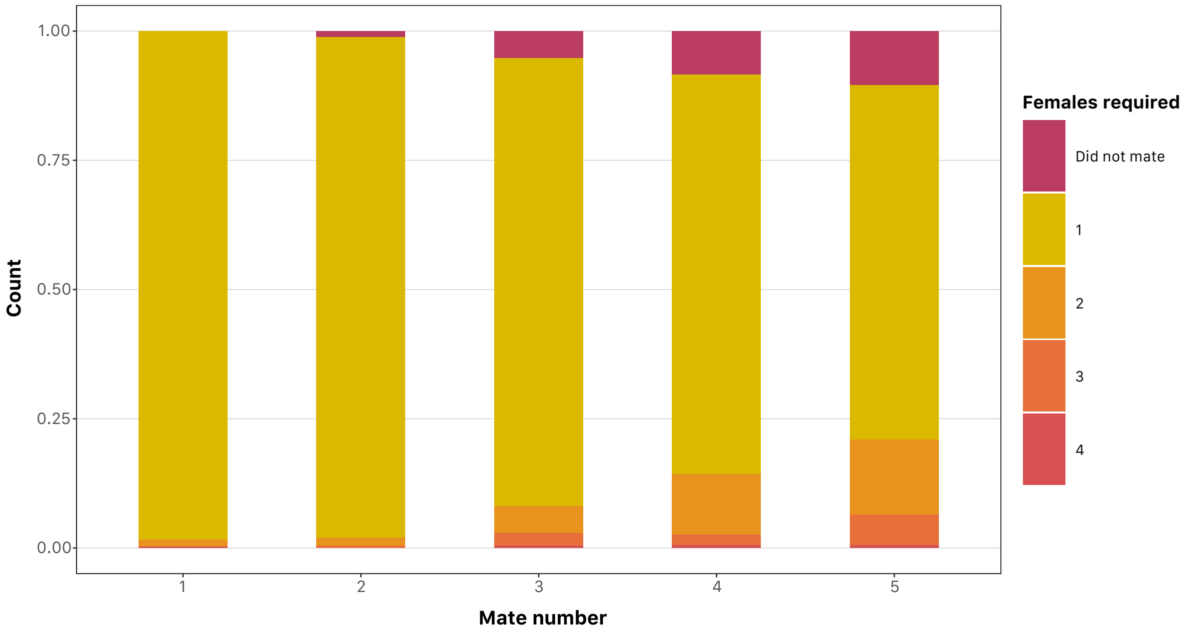


Supplementary Figure 4

The number of virgin females presented to an experimental male before a successful first, second, third, fourth or fifth mating for that male took place. New, additional virgin females were added to mating arenas containing a single male when latency to mate with the previous female was >60 minutes. The "did not mate" category represents males that refused to mate within the period of the mating assay.

| Essential amino acid stock | |
|---|---|
| | (g/200 ml) |
| F (L-phenylalanine) | 3.03 |
| H (L-histidine | 2.24 |
| K (L-lysine) | 5.74 |
| M (L-methionine) | 1.12 |
| R (L-arginine) | 4.70 |
| T (L-threonine) | 4.28 |
| V (L-valine) | 4.42 |
| W (L-tryptophan) | 1.45 |
| **Non-essential amino acid stock** | |
| | (g/200 ml) |
| A (L-alanine) | 5.25 |
| D (L-aspartate) | 2.78 |
| G (glycine) | 3.58 |
| N (L-asparagine) | 2.78 |
| P (L-proline) | 1.86 |
| Q (L-glutamine) | 6.02 |
| S (L-serine) | 2.51 |

Supplementary Table 1

Recipe for essential amino acid and non-essential amino acid stock solutions (Camus et al., 2018).

| Protein diet | | | Carbohydrate diet | |
|---|---|---|---|---|
| L-ile | Powder | 348mg | Sucrose | 6.5g |
| L-leu | Powder | 492mg | | |
| L-tyr | Powder | 252mg | | |
| All following components are identical for both diets | | | | |

| | | |
|---|---|---|
| cholesterol | 20mg/ml in EtOH | 3ml |
| CaCl2 | 1000x | 200µl |
| MgSO4 | 1000x | 200µl |
| CuSO4 | 1000x | 200µl |
| FeSO4 | 1000x | 200µl |
| MnCl2 | 1000x | 200µl |
| ZnSO4 | 1000x | 200µl |
| H2O | | Up to 50ml |
| Autoclave resulting 50ml solutions at this stage | | |
| buffer | 10x acetate buffer base | 20ml |
| Nucleic acid/lipid solution | 125x stock | 1.6ml |
| Essential amino acid solution | Stock (table S1) | 18.154ml |
| Non-essential amino acid solution | Stock (table S1) | 18.154ml |
| Na glutamate solution | 100mg/ml | 5.464ml |
| Cys solution | 50mg/ml | 1.584ml |
| Vitamin mix | 47.6x stock | 4.2 ml |
| Folic acid | 1000x stock | 200µl |
| Propionic acid | | 1.2ml |
| Nipagin | 110g/l stock in 95% EtOH | 3ml |
| Make each to total volumes of 200ml with H2O and syringe filter into tubes for storage<br>Gently warm the protein diet to aid dissolution<br>Add 20% yeast solution to protein diet (at 20% concentration) before use | | |

Supplementary Table 2

Recipe to make 200ml stocks of protein and carbohydrate diet solutions for *Drosophila* (Piper *et al.*, 2014; Camus *et al.*, 2017, 2018). Sucrose in carbohydrate diet is replaced 1:1 by amino acids in the protein diet. Ingredients for the vitamin mix can be found in Piper *et al.*, (2014).

<table>
<tr><th>Data</th><th>Model</th></tr>
<tr><td colspan="2">Intake data</td></tr>
<tr><td rowspan="2">Angle</td><td>glm(alpha ~ block, data=tidy.alpha)</td></tr>
<tr><td>glmmTMB(resid ~ treatment*day + (1 | id), data=tidy.alpha2)</td></tr>
<tr><td rowspan="2">Day 1 angle</td><td>glm(alpha ~ block, data=alpha1)</td></tr>
<tr><td>glmmTMB(resid ~ treatment + (1|id), data=alpha1)</td></tr>
<tr><td rowspan="2">Length</td><td>glm(distance ~ block, data=tidy.distances)</td></tr>
<tr><td>lmer(resid ~ treatment*day + (1 | id), tidy.distances2)</td></tr>
<tr><td rowspan="2">Day 1 length</td><td>glm(distance ~ block, distances1)</td></tr>
<tr><td>glmmTMB(resid ~ treatment + (1|id), data=distances1)</td></tr>
<tr><td rowspan="2">Raw protein</td><td>glm(P ~ block, data=tidyCP, family = Gamma())</td></tr>
<tr><td>glmmTMB(resid ~ treatment*day + (1 | id), data=tidyCPP)</td></tr>
<tr><td rowspan="2">Raw carbohydrate</td><td>glm(C ~ block, data=tidyCP)</td></tr>
<tr><td>glmmTMB(resid ~ treatment*day + (1 | id), data=tidyCPC)</td></tr>
<tr><td>MANOVA</td><td>manova(cbind(alp, dis) ~ treatment*day, data = tidy.alphdis2)</td></tr>
<tr><td colspan="2">Offspring data</td></tr>
<tr><td rowspan="2">Male output</td><td>glm(totaloffspring ~ block, data=offspring2)</td></tr>
<tr><td>glm(resid ~ male, data=offspring2)</td></tr>
<tr><td rowspan="3">Offspring slopes</td><td>c1 %>%group_by(id.b) %>% summarize(slope = coef(lm(adultperday ~ vial))[[2]], .groups = "drop")</td></tr>
<tr><td>glm(slope ~ block, data=coefs)</td></tr>
<tr><td>glmmTMB(resid ~ mate + ( 1 | male), data=coefs)</td></tr>
<tr><td colspan="2">Mating data</td></tr>
<tr><td rowspan="2">Latency, survival</td><td>coxph(Surv(latency, lat.censor) ~ matenumber + block, data = Lmating.times)</td></tr>
<tr><td>survfit(Surv(latency, lat.censor) ~ matenumber, data = Lmating.times)</td></tr>
<tr><td>Latency repeatability</td><td>rptGaussian(latency ~ matenumber + (1 | id), grname = c("id", "Fixed"), data = REPmating.times, nboot = 1000, npermut = 0, adjusted = FALSE)</td></tr>
<tr><td rowspan="2">Duration</td><td>glm(duration ~ block, data=duration.times.2, family = poisson())</td></tr>
<tr><td>glmmTMB(resid ~ matenumber + (1 | id), data=duration.times.2)</td></tr>
<tr><td>Duration repeatability</td><td>rptGaussian(duration ~ matenumber + (1 | id), grname = c("id", "Fixed"), data = REPmating.times, nboot = 1000, npermut = 0, adjusted = FALSE)</td></tr>
<tr><td>Offspring x mating</td><td>glmmTMB(totaloffspring ~ scaledduration + scaledlatency + matenumber + (1|id.b))</td></tr>
</table>

Supplementary Table 3
Statistical models used to analyse data in R version 4.0.4 (The R Foundation for Statistical Computing, Vienna, Austria, http://www.r-project.org).

REFERENCES

Camus, M.F. et al. (2017) ‘Sex and genotype effects on nutrient-dependent fitness landscapes in Drosophila melanogaster’, *Proceedings of the Royal Society B: Biological Sciences*, 284(1869), p. 20172237. Available at: https://doi.org/10.1098/rspb.2017.2237.

Camus, M.F. et al. (2018) ‘Dietary choices are influenced by genotype, mating status, and sex in Drosophila melanogaster’, *Ecology and Evolution*, 8(11), pp. 5385–5393. Available at: https://doi.org/10.1002/ece3.4055.

Piper, M.D.W. et al. (2014) ‘A holidic medium for Drosophila melanogaster’, *Nature Methods*, 11(1), pp. 100–105. Available at: https://doi.org/10.1038/nmeth.2731.
